# Topological defects govern mesenchymal condensations, offering a morphology-based tool to predict cartilage differentiation

**DOI:** 10.1101/2022.05.30.493944

**Authors:** Ekta Makhija, Yang Zheng, Jiahao Wang, Han Ren Leong, Rashidah Binte Othman, Ee Xien Ng, Eng Hin Lee, Lisa Tucker Kellogg, Yie Hou Lee, Hanry Yu, Zhiyong Poon, Krystyn Joy Van Vliet

## Abstract

A critical initial stage of skeletal morphogenesis involves formation of highly compact aggregates of mesenchymal cells, known as mesenchymal condensations, appearing as regularly-spaced pattern of spots. Conventional computational models to understand their patterning have been based on chemotaxis, haptotaxis, and reaction-diffusion equations. In this work, we investigate the mesenchymal condensations from a different perspective, namely topological defects within liquid crystal-like pattern. Using bone marrow-derived mesenchymal stromal cells (bm-MSCs), we observed emergence of cellular swirls in confluent in-vitro cultures, followed by appearance of mesenchymal condensations at the centers of the self-assembled swirls. Specifically, the condensations appeared at the ‘comet-like’ (+1/2) and ‘spiral-shaped’ (+1) topological defect sites within the swirl pattern. Next, with the rationale that cellular swirls precede skeletal morphogenesis, and supported with the qualitative observation that swirl pattern-features are donor-specific, we probed the correlation between swirl pattern and the chondrogenic differentiation outcome of bm-MSCs. Towards this, we first generated and imaged cellular swirls systematically across 5 donors by controlling seeding density, culture vessel geometry, and culture duration. We observed that the swirl pattern features quantified as variance of coherency correlated strongly with the cartilage matrix proteins, sulfated glycosaminoglycan and collagen-II, quantified from the standard *in-vitro* chondrogenic differentiation assay. Our work shows that swirl-pattern quantification provides a novel and powerful tool to predict efficacy of bm-MSCs for *in-vitro* cartilage regeneration.

**Significance Statement:** Mesenchymal condensation is a critical stage in the formation of bone and cartilage, where the mesenchymal cells form high density cell clusters that are regularly spaced. In this work, we inspect the patterning of these condensations in-vitro from a novel perspective. We first show that at high density, bone-marrow-derived mesenchymal stromal cells (bm-MSCs) self-assemble to form cellular swirls resembling the vortices in a turbulent flow. This is followed by cell aggregations at the centers of the vortices, which show correspondence to mesenchymal condensations. Interestingly, we observed that the swirl pattern made by bm-MSCs isolated from human donors, varies from individual to individual and correlates with their propensity to differentiate into cartilage. This suggests that swirl pattern quantification via image analysis can be used to predict differentiation outcome, in context of regenerative cell therapy.

## Introduction

A critical step during the development of skeletal tissues is the self-aggregation of mesenchymal cells, known as mesenchymal condensation [1]–[3]. These cellular condensations appear as a regularly-spaced pattern of spots that correspond to nodules of the developing limb. Conventionally, computational biologists have modelled the patterning of these condensations via chemotaxis, haptotaxis, and reaction-diffusion models [4]–[6]. Recently, there has been growing evidence that patterns in biological systems, such as the cellular swirls that emerge in cell populations, resemble a nematic liquid crystal pattern. In this work, we investigate the mesenchymal condensation patterning from this new perspective.

Cellular swirls that emerge in cell monolayers have been extensively studied in the field of active matter, and modelled as kinetic phase transition [7], jamming transition [8], and glass transition [9]. Recently, these cellular swirls have also been modelled as turbulence in the fluidlike behavior of collective cell motion [10], [11], and as nematic liquid crystal structures with topological defects [12], [13]. Currently, there is excitement in the field [14], [15] with discoveries that the so-called topological defect sites are the scenes for fundamental morphogenetic events, such as epithelial cell extrusion [16] and neural crest cell migration [17]. In this work, we show that mesenchymal condensations form at topological defect sites, followed by its implications in predicting the differentiation efficacy for regenerative cell therapy.

The growing promise of cell therapy for tissue regeneration calls forth a need for critical quality attributes predictive of the potential of cell product to differentiate into desired tissue type. One such example considered here, is MSC therapy for cartilage repair, as cartilage located in joints commonly gets damaged, often resulting in osteoarthritis. The currently used *in-vitro* assay for predicting efficacy of MSCs for cartilage regeneration is the chondrogenic differentiation assay, a relatively slow process of 21 days followed by protein quantification assays [18]. In this work, we propose quantification of swirl pattern in MSC monolayers for early prediction of differentiation efficacy, with the potential of incorporating it as a label-free attribute used in monitoring assay during cell manufacturing process. We believe that swirl pattern being a collective-cell morphological feature would offer a holistic prediction of differentiation potential compared to prediction models based on single-cell morphometric features.

## Results

### Cellular swirls emerge in confluent cultures of bone marrow-derived MSCs (bm-MSCs)

A visual inspection of *in-vitro* cultures of bm-MSCs at confluency showed cellular swirls formed by alignment between neighboring cells (Fig. 1A). To observe the origin of these emergent swirls, time-lapse imaging was performed for 2 weeks, starting from sparsely seeded single cells (Movie S1, Movie S2). We observed that when local cell density reached a critical value, around day 10, the cells transitioned into a collective motion resembling fluid-like behavior. This collective cell flow produced turbulence-like vortices, building towards the appearance of swirls formed by alignment of neighboring cells.

**Figure 1.**
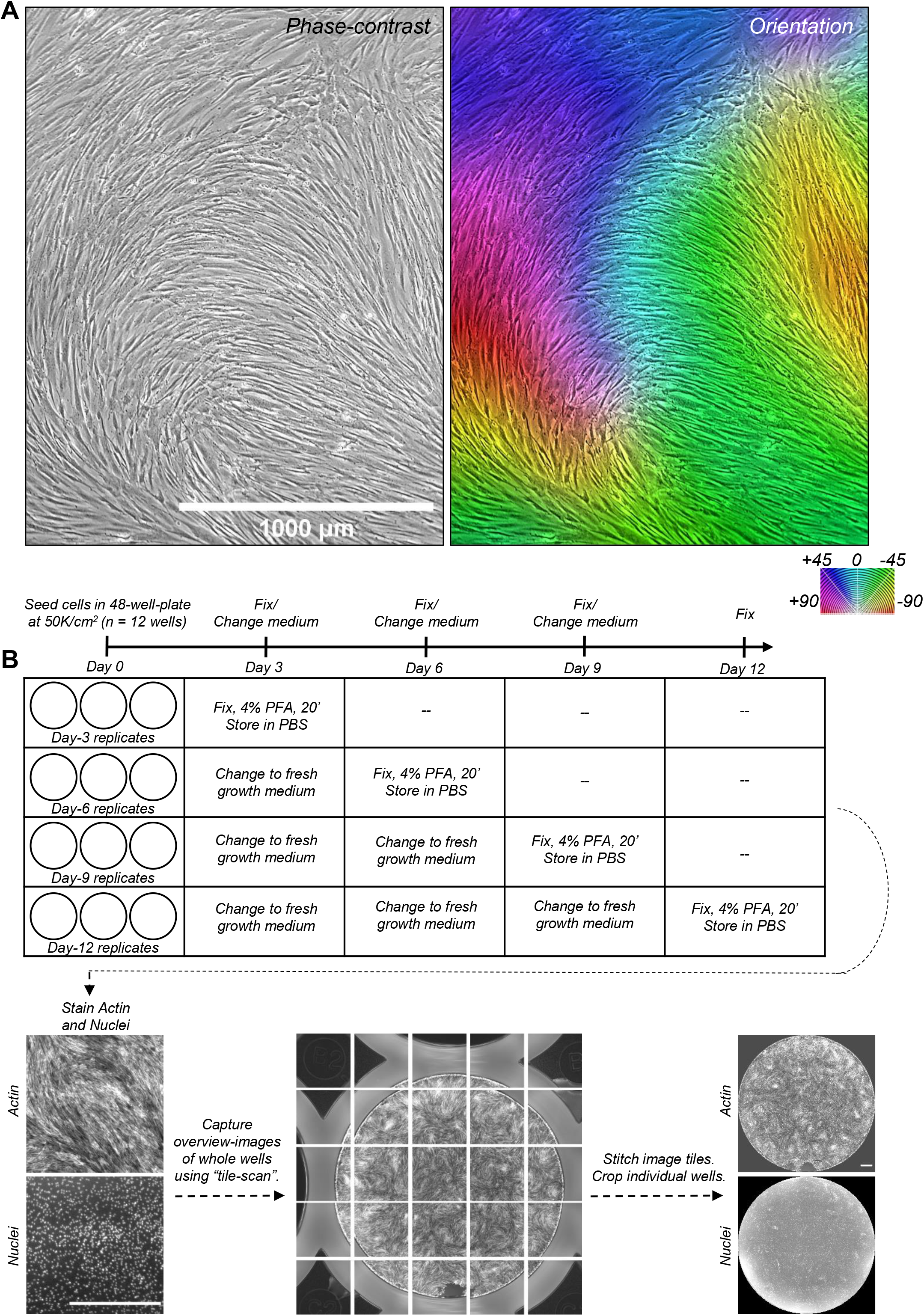
Cellular swirls emerge in confluent cultures of mesenchymal stromal cells. (A) Left panel shows phase-contrast image of bone marrow-derived mesenchymal stromal cells cultured to confluency in T75 flasks. Right panel shows its corresponding orientation image generated using OrientationJ plugin in ImageJ. (B) Experiment methodology showing cell sample preparation, staining, imaging, and image processing to generate whole well pattern images. All scale bars are 1000 μm.

Motivated by the possibility of using the cellular swirl pattern as a morphological feature for prediction of differentiation outcome, we systematically generated and imaged the cellular swirls by controlling initial cell density, geometry of cell culture vessel, and culture duration (Fig. 1B). The value of starting cell density was chosen such that cells are close to 90% confluent upon attachment, as this would minimize the time required for emergence of cellular swirls. The culture vessel chosen was 48-well plate, so as to optimize between the number of swirl features per well and the byte-size of the whole-well overview image. Since the 48-well plates are not suitable for phase contrast imaging, actin and nuclei were stained using phalloidin and DAPI fluorescent stains respectively. Tiled images of actin and nuclei were acquired and stitched to capture the complete pattern formation in the whole well, followed by cropping of individual circular wells. Samples were fixed on day 3, 6, 9, and 12 post seeding. Before proceeding with the quantification of swirl pattern, we closely examined certain features in the pattern, especially the actin arrangement and cell density at centers of the swirls.

### Cells self-aggregate at swirl centers, the so-called ‘positive’ topological defects

The cellular swirls emergent in monolayers have been modelled as nematic liquid crystals [13]. A salient feature of the nematic liquid crystal pattern is topological defects, i.e. the sites within the pattern where sudden changes in orientation occur, for example, the centers of the spiral swirls [19]. Zoomed-in images of ‘comet-like’ (+1/2) and ‘spiral-shaped’ (+1) topological defect sites, in MSC swirl pattern, show extremely high cell density clusters (often with overlapping nuclei, Fig. 2, Fig. S1), while the −1/2 type topological defect sites show local minima of cell density (Fig. S2). Such clustering can be induced by turbulence-like flow in active systems as is recently being explored [20]. These high density cell clusters that appear at swirl centers resemble mesenchymal condensations – the self-aggregation of mesenchymal cells during early stage of cartilage and bone development [1]–[3].

**Figure 2.**
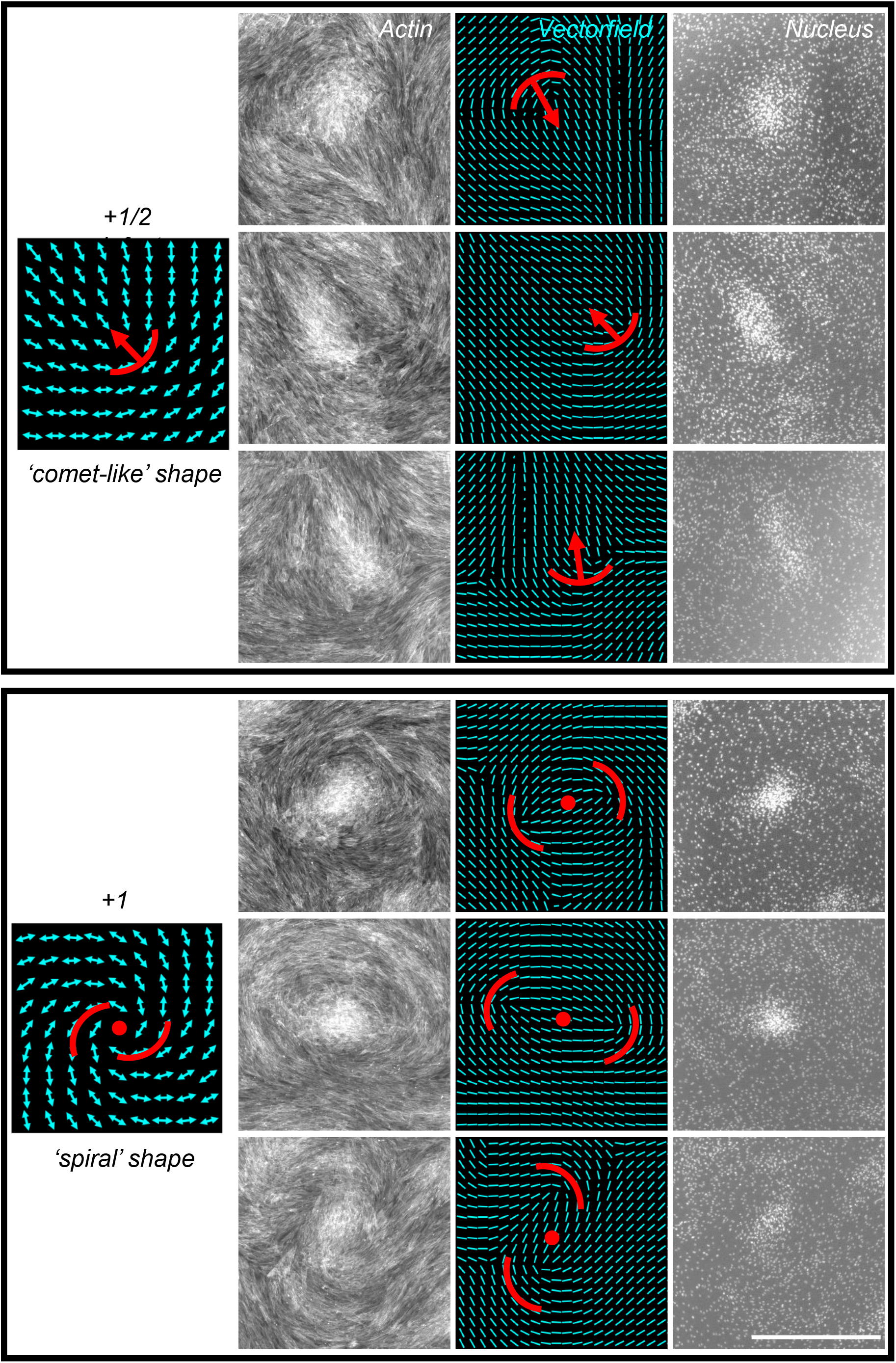
Cells aggregate at +1/2 and +1 topological defect sites. Six regions (cropped from whole-well stitched images) showing that cells aggregate at sites of +1/2 and +1 topological defects. The aggregates are visible in the nucleus images, while the defects are visible in the actin and corresponding orientation vectorfield images. Orientation vectorfields were generated using the OrientationJ plugin in ImageJ (see Methods). Scale bar, 1000 μm.

### Self-aggregated cell clusters correspond to mesenchymal condensations

To test whether the cells that aggregate at +1/2 and +1 defect sites indeed correspond to mesenchymal condensations, we stained the day 8 samples with fluorescent tagged peanut agglutinin (PNA) antibody, a lectin binding protein used for labelling mesenchymal condensations [21]. Zoomed in images show high intensity of PNA at the cell clusters (Fig. 3). This suggests that the mesenchymal condensations observed in *in-vitro* cell cultures arise from the density fluctuations caused by the self-assembled cellular swirls. Further, immunostaining of transcription factors YAP, SOX9, and RUNX2 showed high intensity in the nuclei within the cell aggregates (Fig. S3). This suggests a possibility that the high mechanical stresses, known to arise at +1/2 and +1 defect sites, activate the mechanosensitive transcription factor YAP in the mesenchymal condensations, followed by downstream activation of RUNX2 and SOX9, the master transcription factors of osteogenic and chondrogenic differentiation [22].

**Figure 3.**
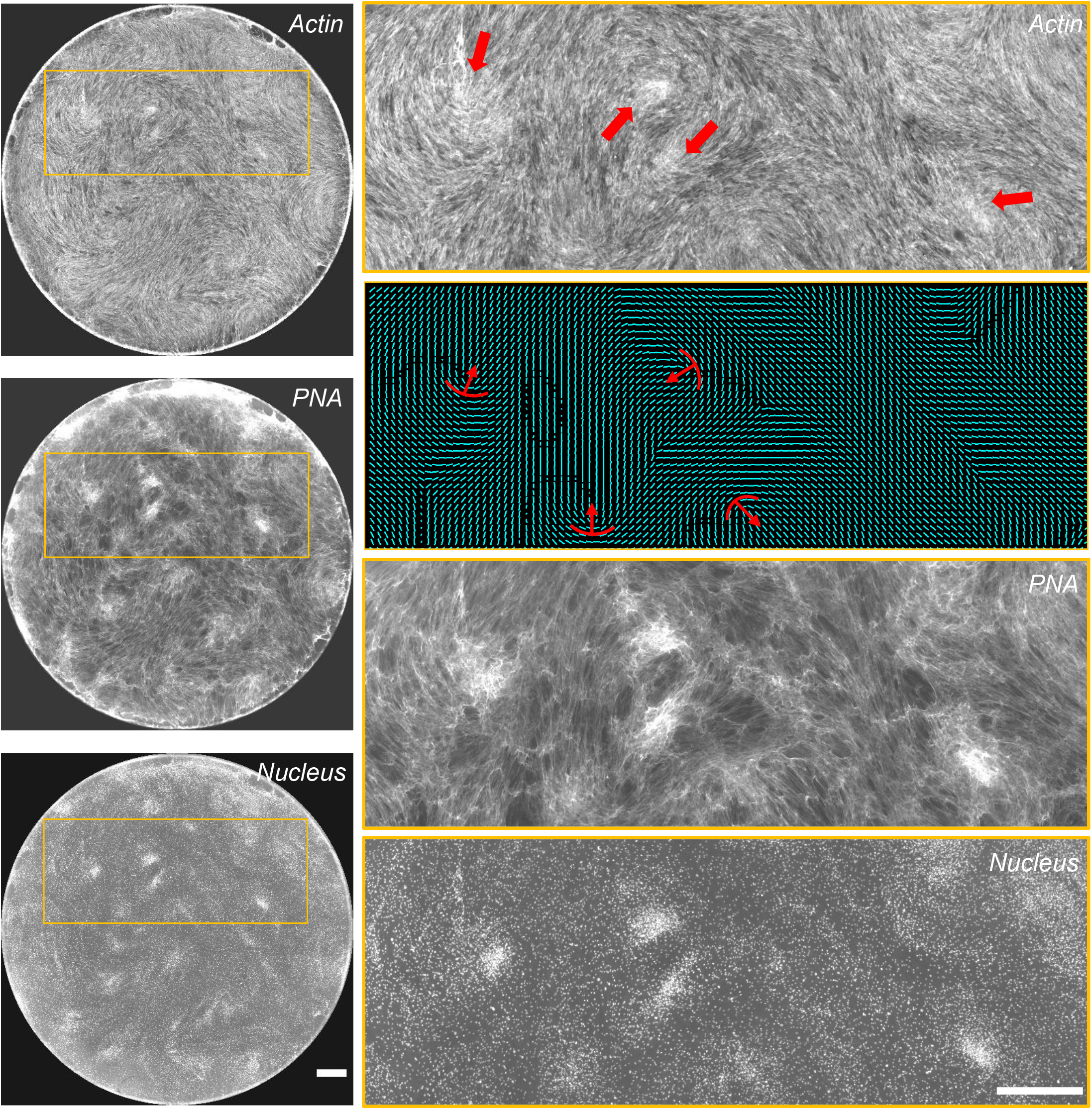
Cell aggregates at topological defects correspond to mesenchymal condensations. The cell aggregates arising at defect sites of actin nematic pattern colocalize with peanut agglutinin (PNA), the marker for mesenchymal condensations - a pre-requisite for formation of cartilage or bone during skeletal development. Scalebar 1000 μm

### Swirl pattern is donor-specific and reproducible

We first performed a qualitative comparison of the swirl pattern made by bm-MSCs across multiple donors, via phase contrast images from cultures maintained by different researchers (without intentional control of seeding density, geometry of culture vessel, and culture duration) (Fig. 4A). This comparison, as well as comparison of swirl pattern from actin images of systematically generated cellular swirls in 48-wells (Fig. 4B), showed that swirl pattern features are donor-specific and reproducible across replicates. Next, we tested whether a quantification of this swirl pattern across various donors correlates with their chondrogenic differentiation outcome, since cellular swirls lead to mesenchymal condensations, a crucial step that precedes chondrogenic differentiation.

**Figure 4.**
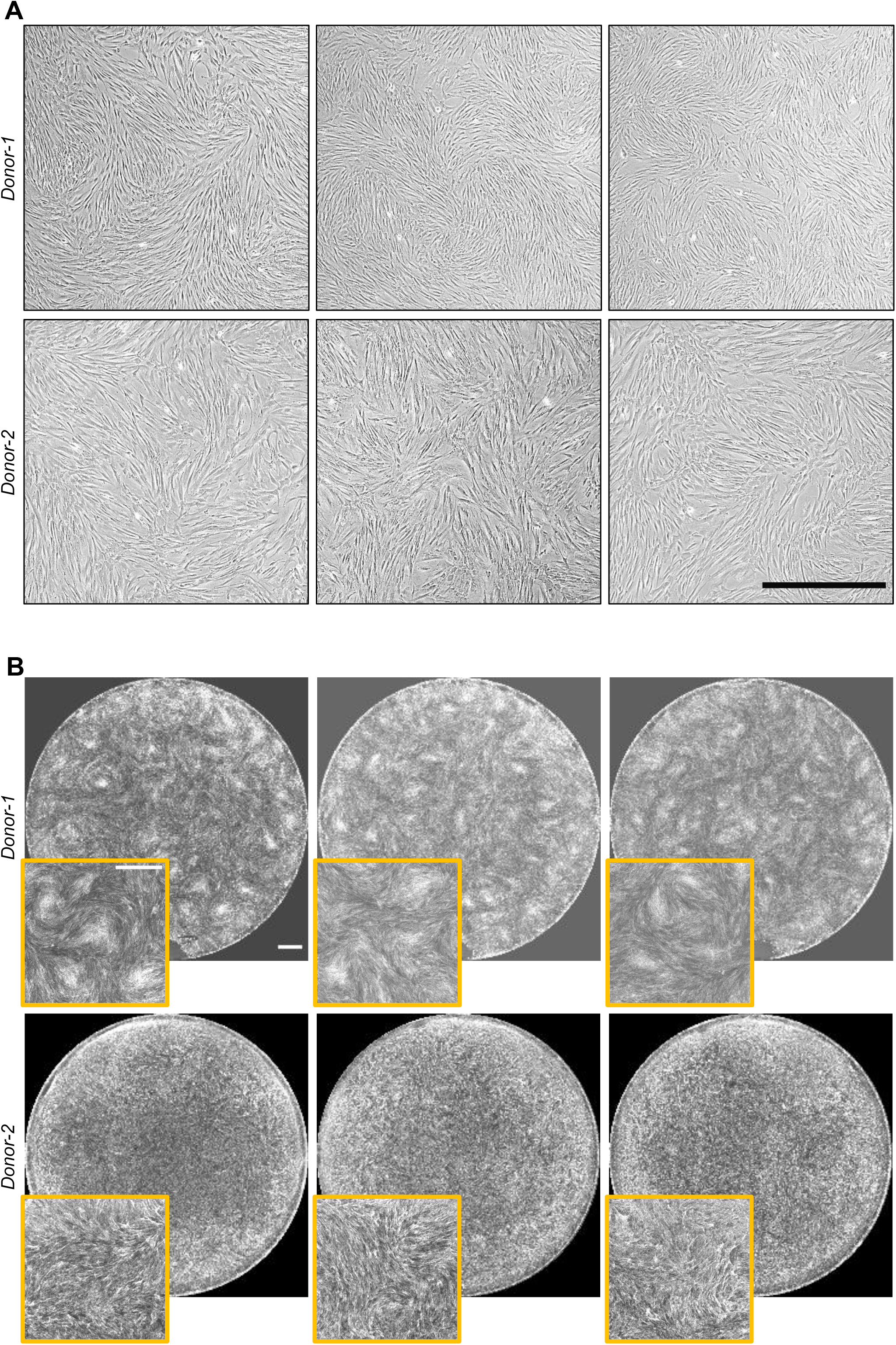
Self-assembled cellular swirl pattern is donor-specific and reproducible. (A) Phase-contrast images at confluency of bm-MSCs from 2 different donors cultured by different researchers (without intentional control of seeding density, geometry of culture vessel, and culture duration). (B) Whole-well actin images of systematically generated cellular swirls on day 3 post-seeding. The bottom-panel shows the regions cropped from the whole-well actin pattern. All scale bars 1000 μm

### Swirl pattern quantification correlates with chondrogenic differentiation outcome

To test this correlation between swirl pattern and differentiation outcome, first, the chondrogenic differentiation assay was conducted for 5 donors (see Methods). Briefly, this assay involved 3 week pellet culture in chondrogenic differentiation condition, followed by measurement of expression levels of cartilage matrix components sulfated glycosaminoglycans (sGAGs) and collagen-II (Col2). Simultaneously, cellular swirls were generated and imaged for the same 5 donors at various time points post cell seeding on 48-well plates. For the quantification of swirl pattern, two different methods were adopted; one using nucleus image, and the other using actin image. In the first method, the total area of cell clusters was quantified, for which the nucleus images were thresholded, and the total area of segmented regions was measured (Fig. S4A, S4B, see Methods). While the total area of clusters per well increased from day 3 to day 12 for all donors (Fig. S4C), it did not correlate strongly with the cartilage matrix protein levels (Pearson’s correlation coefficient, r < 0.5). This could have two possible implications: (1) higher total area of clusters does not necessarily mean better chondrogenesis, or (2) the total area computed from intensity-based thresholding does not successfully capture the quality of mesenchymal condensations. Since an alternate thresholding based on either the size of the cluster or the cell density within the cluster was non-trivial, another method to quantify swirl pattern from actin images was adopted. The second method was based on the reasoning that condensations occur only in the region where the pattern has a single large defect as opposed to regions with short-range order (with multiple small defects) or long-range order (without defects). For the purpose of illustration, we cropped from whole-well actin images, three such regions corresponding to each type of pattern (Fig. 5A). The correspondingly cropped nucleus images showed occurrence of condensation only for the large defect region. Next, the orientation and coherency images were generated from the three actin images (see Methods). The orientation, which shows the direction of local features, varies from −90 to +90 degrees. The coherency, which measures the local structure, is 1 when the local structure has a dominant orientation, and 0 when the local structure is unaligned or isotopic. Subsequently, the variance of coherency (VoC) was computed for the three coherency images, which showed that the large defect region has higher VoC compared to regions with either short-range order (with multiple small defects) or long-range order (without defects).

**Figure 5.**
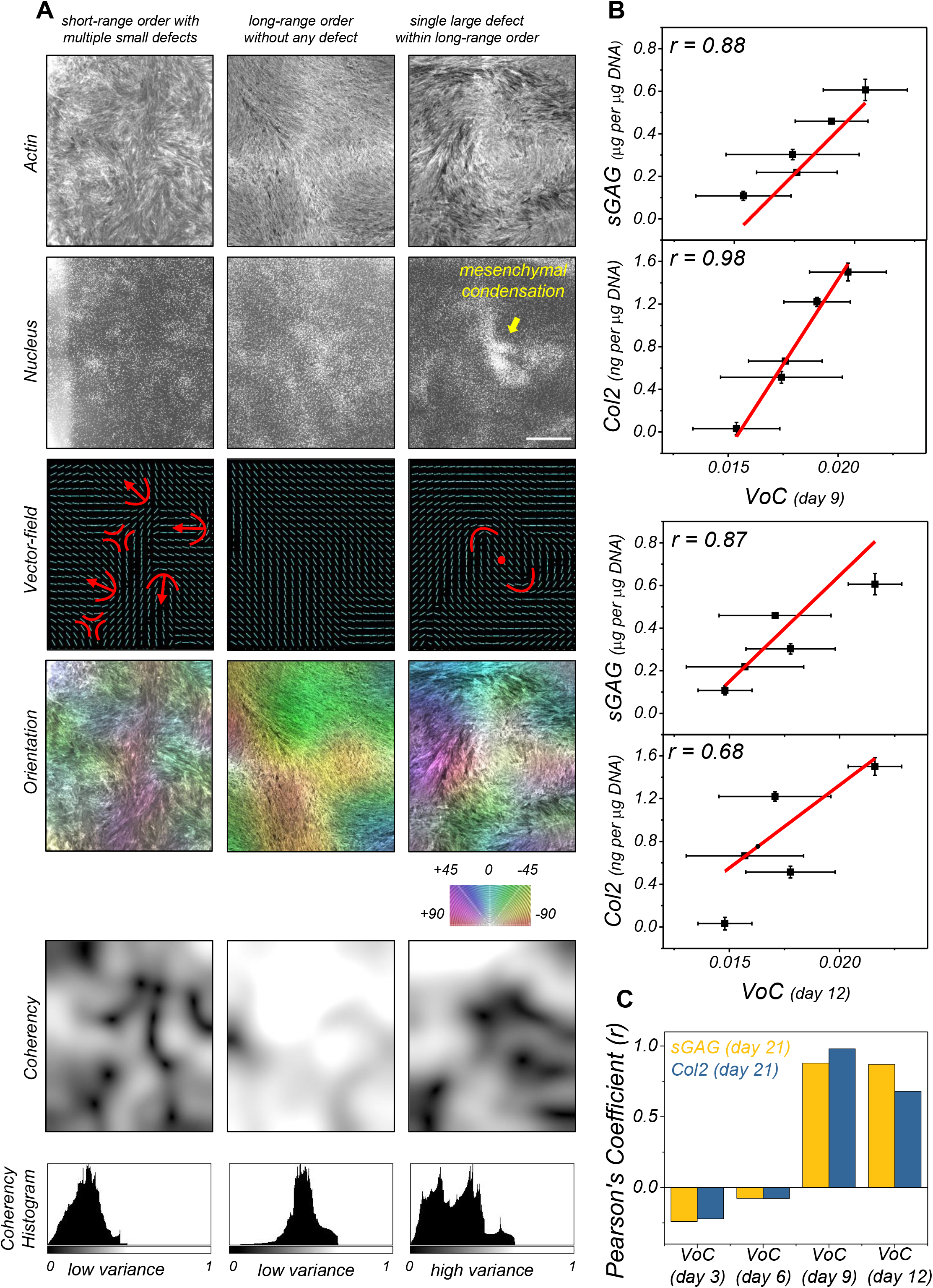
Variance of pattern coherency correlates with chondrogenic differentiation. (A) The first and second rows show nucleus and actin stain in three regions where the pattern may be classified as having ‘*short-range order*’, ‘*long-range order*’, and ‘*topological defect*’. *The t*hird, fourth and fifth rows show vectorfield, orientation and coherency images for the three regions, generated from the above actin images using OrientationJ (see Methods). The values of orientation vary from −90 to +90 degrees, and the values of coherency vary from 0 to 1. The last row shows histogram of coherency images (B) Scatter plots of day 9 and day 12 variance of coherency (VoC) vs cartilage matrix proteins sGAG and Col2 quantified from the 21 day chondrogenesis differentiation assay. Red line shows the linear fitting. (C) Pearson correlation coefficient for day 3, 6, 9, 12 VoC vs levels of matrix protein, n = 5 donors. Scale bar 1000 μm

Following this method, VoC was computed from all whole-well actin images (5 donors, 4 time points, 3 replicates, see Methods). Interestingly, the VoC at day 9 and 12 showed strong correlation with both cartilage matrix components, generating a Pearson’s correlation coefficient, r > 0.85 for sGAG, and r > 0.68 for Col2 (Fig. 5B, 5C). The low correlation of VoC at earlier time points is possibly because of lag in emergence of cellular swirls across donors, since differences in cell morphology and proliferation rates would affect the value of critical density at which the swirls emerge and the time required to achieve that critical density.

## Discussion

In this work, we systematically generated and analyzed the self-assembled cellular swirls that emerge in confluent bm-MSCs by controlling seeding density, culture vessel geometry, and culture duration. We showed that the cell clusters that self-aggregate at the centers of the swirls, at the so-called ‘comet-like’ (+1/2) and ‘spiral-shaped’ (+1) topological defects, correspond to mesenchymal condensations. We showed that this swirl pattern is donor-specific and reproducible across replicates. We demonstrated implications of swirl pattern in predicting differentiation outcome – the variance of coherency (VoC) of swirl pattern strongly correlates with expression of cartilage matrix components when compared across multiple donors. This implies the possibility of using swirl-pattern quantification as a critical quality attribute for predicting efficacy of mesenchymal stromal cell therapy product for cartilage repair. Further implications of this work for cell therapy manufacturing are discussed below.

A crucial stage during the cell therapy manufacturing is that of cell expansion. Typical protocol optimized for expansion recommends seeding mesenchymal stromal cells sparsely and harvesting them before they reach confluency [23], [24]. Hence, a prediction model based on single-cell morphological features could be easily incorporated in-line within the manufacturing pipeline. While cellular swirls only appear at high confluency, advances in forecasting of active nematics [25] could pave the way for an earlier prediction from low confluency cell culture images. Secondly, with the new understanding that cellular swirls lead to mesenchymal condensations, cell manufacturers for cartilage regeneration could consider culturing MSCs beyond confluency to prime the cells towards chondrogenesis. In such a case, an in-line label-free monitoring of chondrogenic differentiation efficacy via swirl pattern would be feasible, as we have shown that VoC can also be measured from phase-contrast images (Fig. S5). Thirdly, culturing MSCs on engineered patterned surfaces that induce formation of topological defects [26] would provide a better control for manufacturing desired cell therapy product.

Growing evidence from *in-vivo* cartilage repair studies shows that the implanted MSCs play an indirect role in tissue repair via secretion of paracrine factors as opposed to directly differentiating into chondrocytes [27]. While our current work provides an early prediction of outcome 3 week-long *in-vitro* differentiation assay, ongoing experiments in our lab are aimed at comparing this assay with alternate *in-vitro* assays for their ability to predict *in-vivo* cartilage repair outcome.

## Materials and Methods

### 1. Cell-source and cell expansion

Human adult bone marrow-derived MSCs (bm-MSCs) from 5 different donors were purchased from Lonza and StemCell Technology. For the first round of expansion, these cells were seeded in culture flasks (ThermoFisher Nunc) at 2000 cells/cm^2^ using growth medium comprising of Dulbecco’s modified Eagle’s medium, low glucose and pyruvate (Gibco), supplemented with 10% MSC certified fetal bovine serum (Gibco) and 1% penicillin and streptomycin (Gibco). Medium was changed every 3 days until the cells reached ~80% confluency, when the cells were harvested using the trypsin (Gibco). Typically, expanded bm-MSCs were used at passage 3-4.

### 2. Sample preparation for generating cellular swirls

Cells were seeded in 48-well plates at a density of 50,000 cells/cm^2^. Note that the first and last row, as well as the first and last column of the well-plate were not used, because limitations in XY translation of the microscope stage affects the whole-well imaging for these wells. Also, samples corresponding to different time-points were seeded in different well-plates for the reason that paraformaldehyde (PFA) vapors from the samples being fixed resulted in cell death in other samples within the same well plate. For the time-study experiments, the samples were fixed on either day 3, 6, 9, or 12 with medium change every 3 days until they’re fixed. For fixing, growth medium was replaced with 4%PFA in PBS (Biotium, 200ul per well) for 20 minutes, following which the PFA was replaced with PBS. The fixed samples were stored in the incubator or refrigerator until all time points were fixed and ready for staining.

### 3. Fluorescent Staining

Fixed samples were first permeabilized using 0.1% triton X-100 detergent solution (Thermo Scientific) for 5 minutes, followed either by blocking and primary antibody staining or directly by 30 minute staining of nucleus (NucBlue Fixed Cell ReadyProbes, Life Technologies) and actin (ActinGreen 488 ReadyProbes, Life Technologies).

For antibody staining, permeabilization step was followed by an hour of incubation with blocking buffer comprising 1% bovine serum albumin solution (Sigma) with 0.1% tween (Sigma). Next, the primary antibody, diluted in blocking buffer, was added and samples were incubated overnight at 37°C. Peanut agglutinin (lectin PNA from Arachis hypogaea, Alexa Fluor 488 Conjugate, Life Technologies, dilution 1:20), SOX9 (rabbit monoclonal to SOX9 Alexa Fluor 647, Abcam, dilution 1:50), RUNX2 (rabbit monoclonal RUNX2, Alexa Fluor 647, Abcam, dilution 1:50), and YAP (rabbit monoclonal to active YAP1 Alexa Fluor 488, Abcam, dilution 1:50) antibody staining were performed in day 8, day 9, or day 10 samples.

### 4. *In-vitro* chondrogenic differentiation assay

MSCs were centrifuged down into pellet of 1x 10^5^ cell in 96 well non-adhesive plate (Nest Biotechnology Co., Ltd.). Cells were cultured in chondrogenic differentiation medium, which consists of high glucose DMEM supplemented with 10^-7^M dexamethasone, 1% ITS + premix, 50 μg/ml ascorbic acid, 1 mM sodium pyruvate, 0.4 mM proline and 10 ng/ml TGF-β3 (R&D Systems, Minneapolis, MN). Differentiation was carried out for 3 weeks, with medium change every 3 days.

### 5. Sulfated glycosaminoglycan (sGAG) and type II collagen quantification

Samples were digested with 10 mg/mL of pepsin in 0.05 M acetic acid at 4°C, followed by digestion with elastase (1 mg/mL). The amount of sulfated glycosaminoglycan (sGAG) was quantified using Blyscan sGAG assay kit (Biocolor, UK) according to manufacturer’s protocol. Absorbance was measured at 656 nm and sGAG concentration was extrapolated from a standard curve generated using a sGAG standard. Type II Collagen (Col 2) content was measured using a captured enzyme-linked immunosorbent assay (Chondrex, Redmond, WA). Absorbance at 490 nm was measured and the concentration of Col 2 was extrapolated from a standard curve generated using a Col 2 standard. The amount of both sGAG and Col 2 content were normalized to the DNA content of respective samples, measured using Picogreen dsDNA assay (Molecular Probes, OR, USA). Three replicates were analyzed within each group.

### 6. Static imaging and image pre-processing

Whole-well overview images of actin, nucleus, and other fluorescently-labelled proteins (PNA, SOX9, RUNX2, YAP) were captured on Olympus IX83 fluorescence microscope using 4X objective and tile-scan option in Metamorph software. Individual tiles were 2048 x 2048 pixels, pixel size was 1.6 microns. Image tiles were stitched using the “Stitching” plugin in ImageJ [28]. Individual wells were then cropped manually from the stitched images using the circular cropping tool in ImageJ (radius 3636 pixels). Note that the stitching often generated artefact in the form of high intensity at the junctions between neighboring tiles. However, we observed that these intensity artefacts did not affect quantification of coherency from whole-well actin images.

### 7. Quantification from nucleus images

Nucleus images of individual wells were imported into the MATLAB Image Processing Toolbox (Mathworks). First, these images were cropped with a circle mask (radius 2136 pixels) to eliminate the influence caused by uneven light in the corner of wells and then adjusted by contrast enhancement using contrast limited adaptive histogram equalization [29]. For noise removal, two binary masks were generated separately via image global threshold and local adaptive threshold, and then used together to subtract the background. De-noised nucleus images were first processed with a Gaussian filter (radius 100 pixels) to obtain the nucleus distribution heatmap. This heatmap was then normalized and transformed into a binary mask using a threshold of 0.5. Total cell aggregate area per well was estimated from this binary mask.

### 8. Quantification from actin images

Orientation and coherency images were generated from whole-well actin images in ImageJ via the plugin OrientationJ [30], with a local window size of 100 pixels and ‘Gaussian’ gradient-type. Coherency images were cropped manually using the circular cropping tool in ImageJ (radius 3336 pixels) to remove the edge effects. Histogram of the coherency image was generated in ImageJ using Analyze ➔ Histogram. The values of variance were read-out from the histograms.

### 9. Live cell time-lapse imaging

14-day time-lapse imaging to visualize the transition from single cell to cellular swirls was performed as described in an earlier work [31]. Briefly, MSCs were seeded very sparsely at 100 cells / cm^2^ in a glass bottom petri dish, which was mounted on the microscope stage-top incubator. Phase-contrast images were captured using the 10X objective every fifteen minutes. Initially, tile images in a 2×2 grid were captured around the single cells. As the cells migrated and proliferated, the number of tiles was increased to 3×3 on day 4, 4×4 on day 8, and 5×5 on day 12 of the time-lapse imaging.

## Acknowledgements

We thank George Barbastathis, Irmgard Bischofberger, Jongyoon Han, Tetsuya Hiraiwa, and Yusuke Toyama for useful discussions. This research was funded by the National Research Foundation, Prime Minister’s Office, Singapore under its Campus for Research Excellence and Technological Enterprise (CREATE) programme, through Singapore MIT Alliance for Research and Technology (SMART): Critical Analytics for Manufacturing Personalised-Medicine (CAMP) Inter-Disciplinary Research Group.

## Supplementary Figure legends

**Figure S1.**
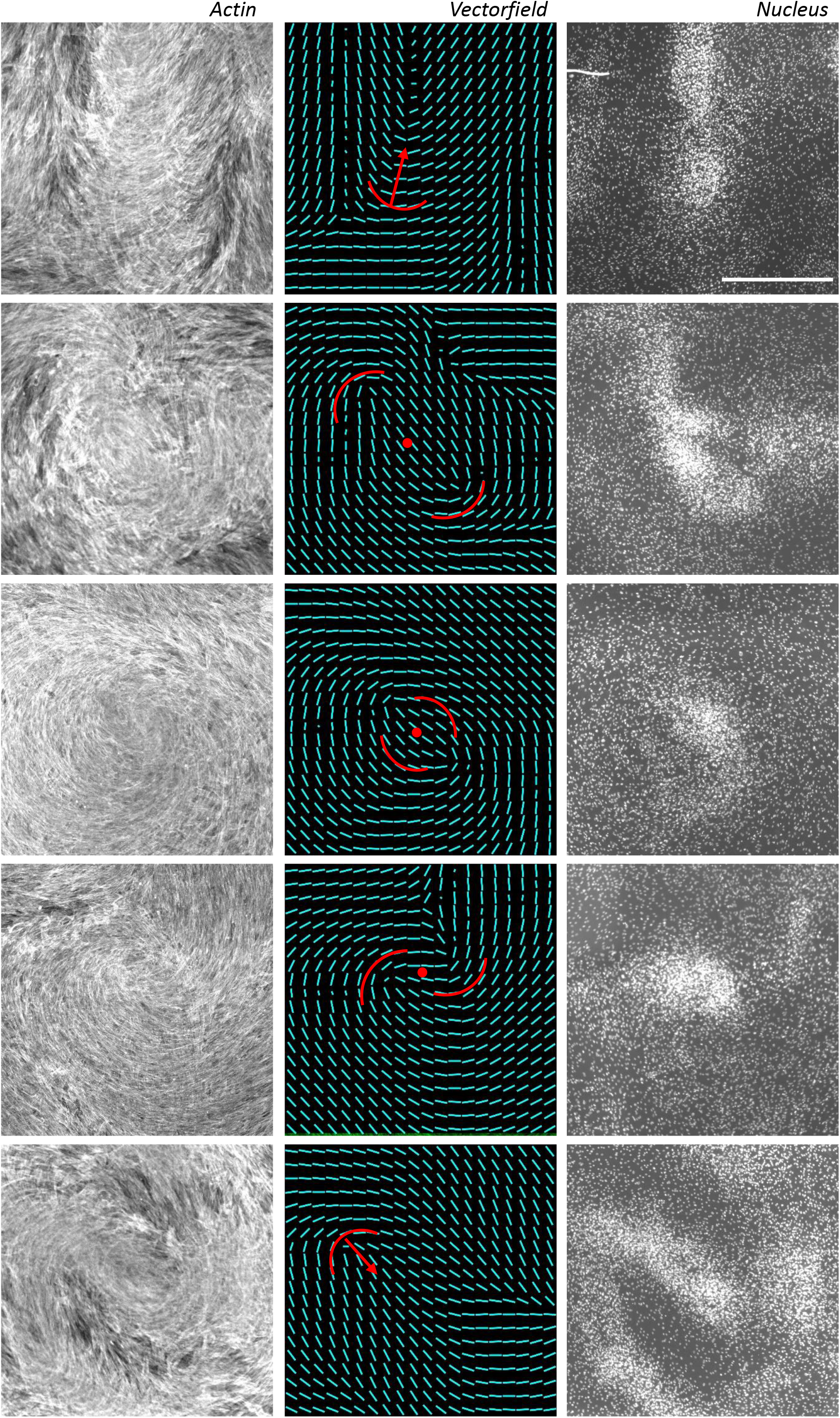
Additional images of cell aggregation at +1/2 and +1 topological defects Five regions (cropped from whole-well stitched images) showing that cells aggregate at sites of +1/2 and +1 topological defects. The aggregates are visible in the nucleus images, while the defects are visible in the actin and corresponding orientation vectorfield images. Orientation vectorfields were generated using the OrientationJ plugin in ImageJ (see Methods). Scale bar 1000 μm

**Figure S2.**
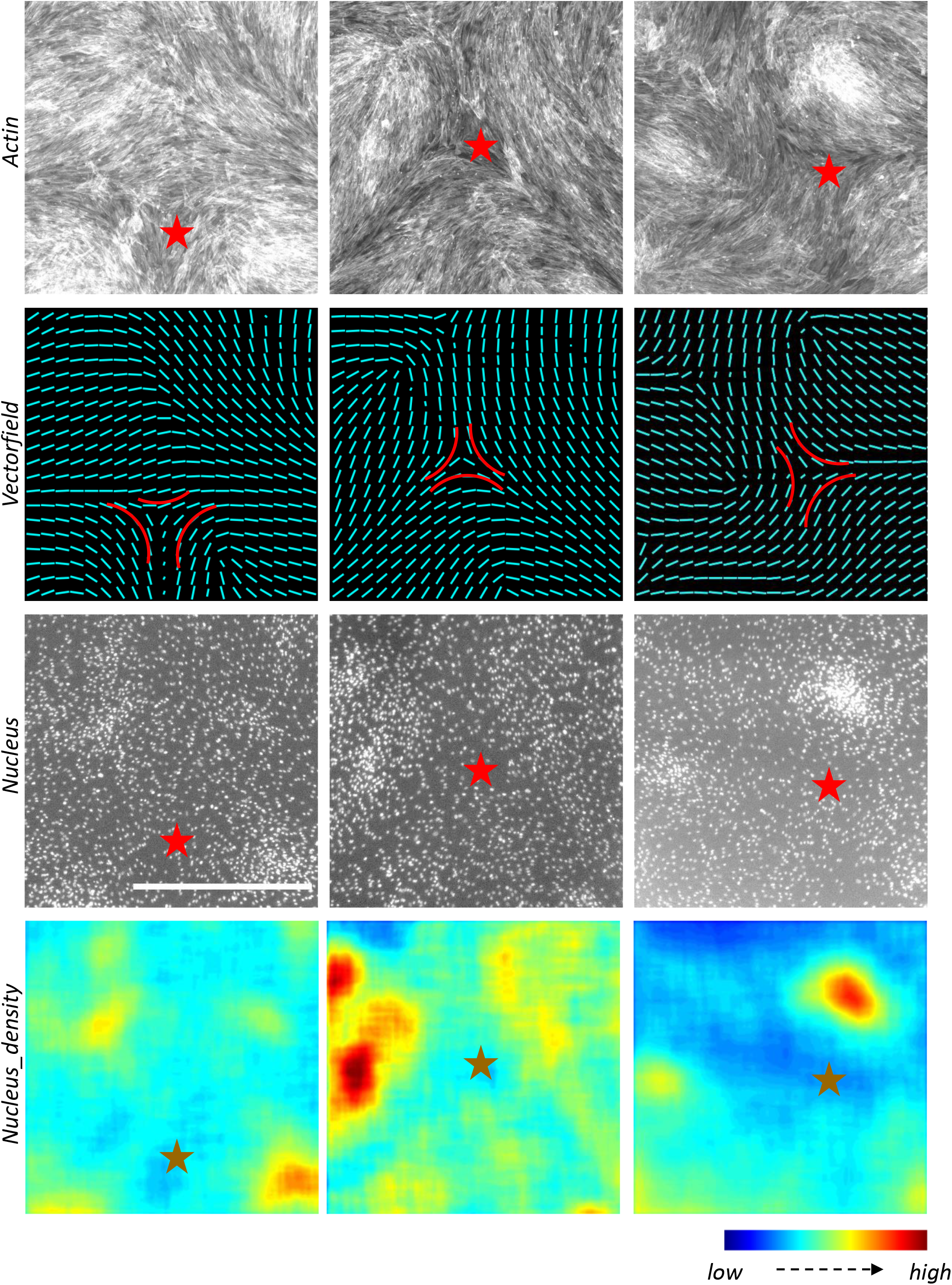
−1/2 topological defect sites have local minimum in cell density. Three regions (cropped from whole-well stitched images) showing that cells recede from −1/2 topological defects. The defects are visible in the actin and corresponding orientation vectorfield images, while the density-minima are visible in the nucleus density colormap. Orientation vectorfields were generated using the OrientationJ plugin in ImageJ (see Methods). Nucleus density colormaps were generated using an averaging filter (100 pixels, i.e., 16 μm) on the nucleus intensity images in MATLAB. Scale bar 1000 μm

**Figure S3.**
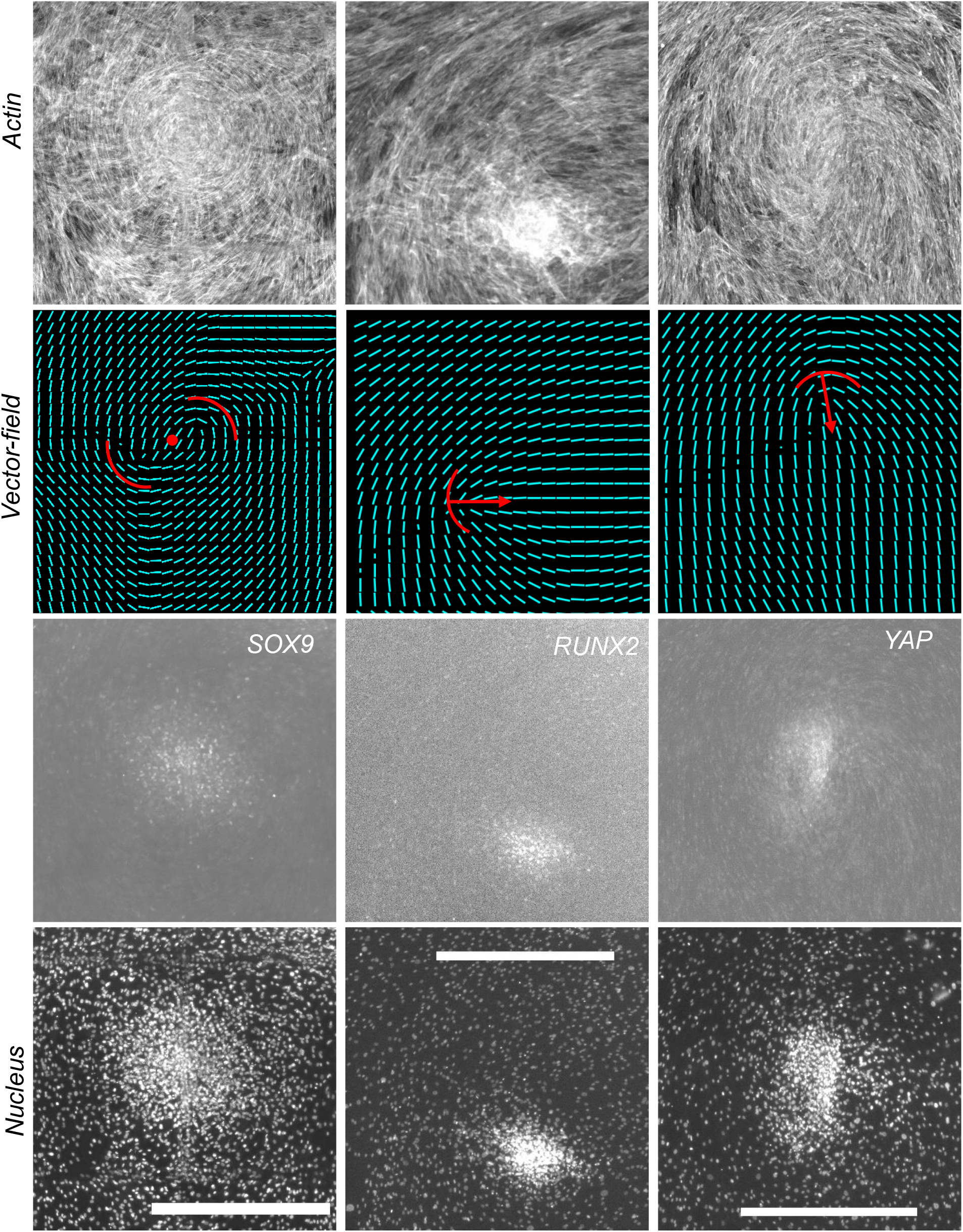
Upregulation of SOX9, RUNX2, and YAP in cell aggregates at defect sites Regions cropped from whole-well stitched actin, nucleus, and transcription factor images show higher levels of SOX9, RUNX2, and YAP in the nuclei of cells aggregated at the topological defect sites. SOX9, RUNX2, and YAP were stained in day 10, day 9, and day 8 samples respectively. Scalebar, 1000 μm

**Figure S4.**
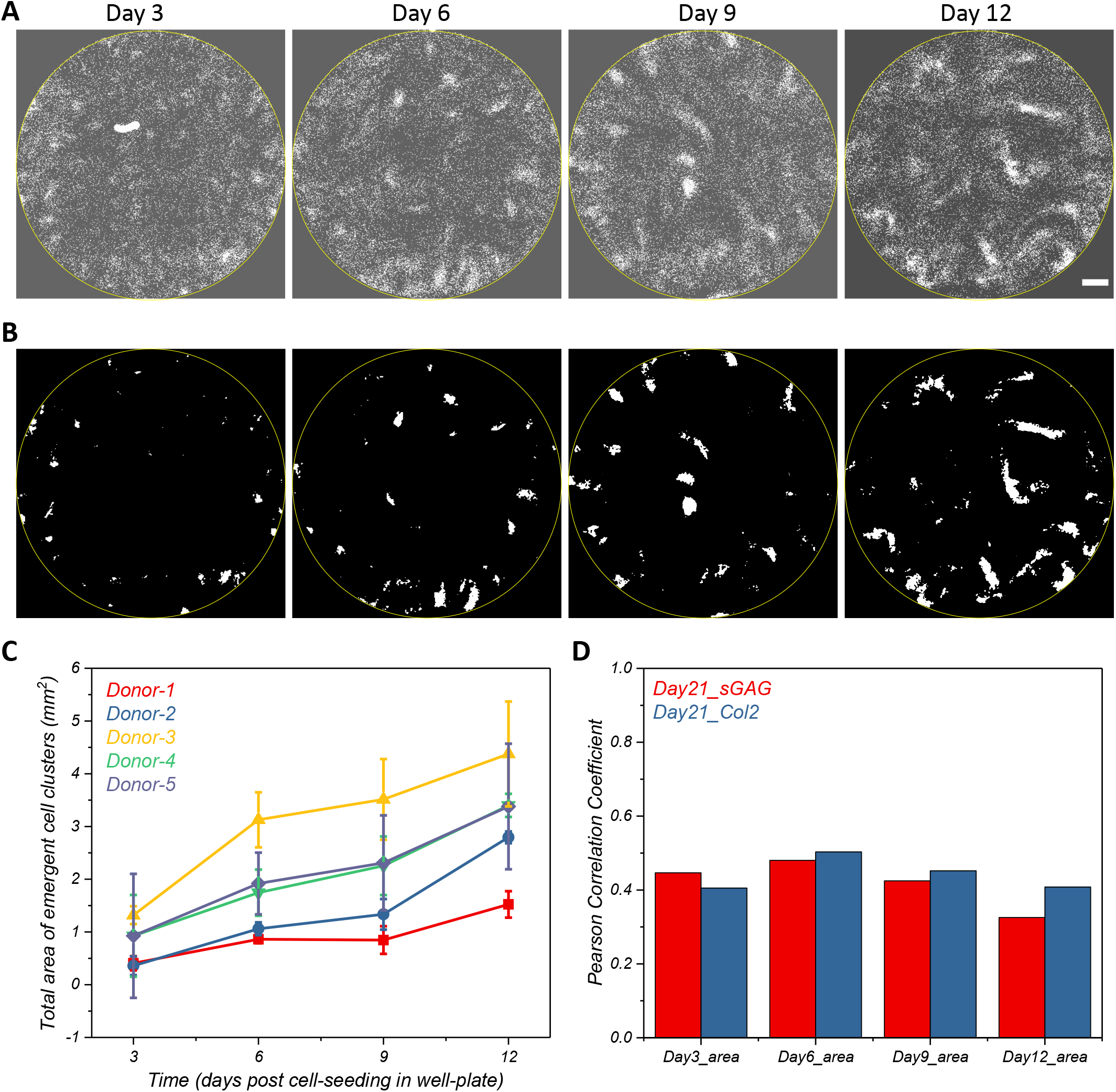
Temporal dynamics of total cell aggregate area (A) Whole-well stitched nuclei images from a single donor at time 3, 6, 9, 12 days post seeding. (B) Binary masks generated by thresholding the above nuclei images. Spots corresponding to debris were manually removed. (C) Total cell aggregate area per well plotted as a function of time for 5 MSC donors (n = 3 replicates per donor). (D) Pearson correlation coefficient for day 3, 6, 9, 12 total area vs levels of matrix protein, n = 5 donors. Scalebar, 1000 μm

**Figure S5.**
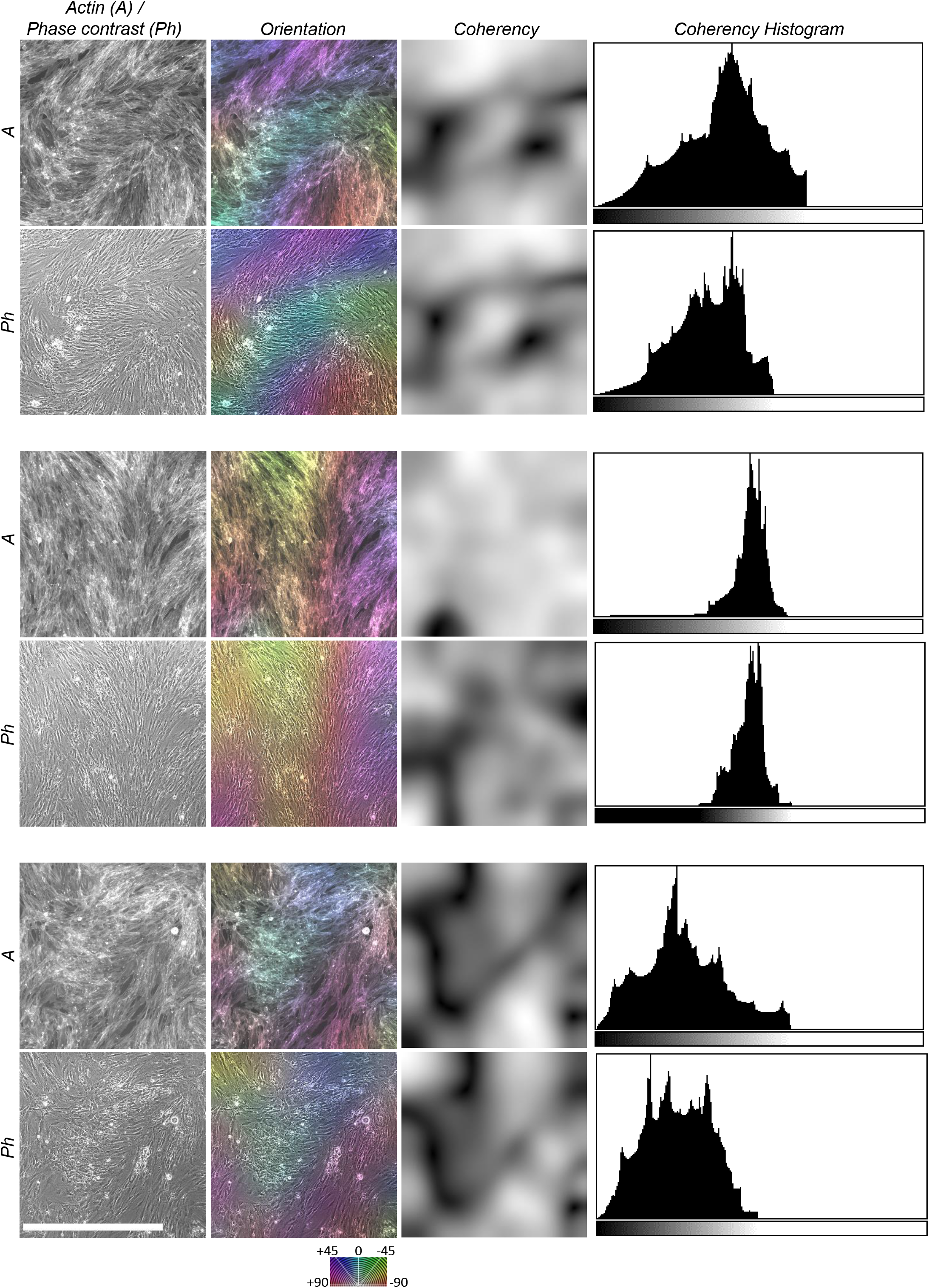
Label-free quantification of orientation and coherency Comparison of orientation analysis for three different regions shows similarity in orientation, coherency and its variance from actin vs phase-contrast images. Scalebar, 1000 μm

## Supplementary movie legend

**Movie S1**. Single cell to cellular swirls

14-day time lapse phase-contrast imaging showing emergence of cellular swirls. Yellow box outlined on the left corresponds to the zoomed region shown on the right.

**Movie S2**. Critical density to cellular swirls

Time lapse of the last 3 days sub-stacked from Movie S1.

## References

[1] B. K. Hall and T. Miyake, “Divide, accumulate, differentiate: Cell condensation in skeletal development revisited,” Int. J. Dev. Biol., vol. 39, no. 6, pp. 881–893, 1995.

[2] B. K. Hall and T. Miyake, “All for one and one for all: Condensations and the initiation of skeletal development,” BioEssays, vol. 22, no. 2, pp. 138–147, 2000.

[3] J. L. Giffin, D. Gaitor, and T. A. Franz-Odendaal, “The forgotten skeletogenic condensations: A comparison of early skeletal development amongst vertebrates,” J. Dev. Biol., vol. 7, no. 4, 2019.

[4] R. A. Rolfe, C. A. Shea, and P. Murphy, “Geometric analysis of chondrogenic self-organisation of embryonic limb bud cells in micromass culture,” Cell Tissue Res., vol. 388, pp. 49–62, 2022.

[5] M. A. Kiskowski et al., “Interplay between activator-inhibitor coupling and cell-matrix adhesion in a cellular automaton model for chondrogenic patterning,” Dev. Biol., vol. 271, no. 2, pp. 372–387, 2004.

[6] S. Christley, M. S. Alber, and S. A. Newman, “Patterns of mesenchymal condensation in a multiscale, discrete stochastic model,” PLoS Comput. Biol., vol. 3, no. 4, pp. 743–753, 2007.

[7] T. Vicsek, A. Czirok, E. Ben-Jacob, I. Cohen, and O. Shochet, “Novel Type of Phase Transition in a System of Self-Driven Particles,” Phys. Rev. Lett., vol. 75, no. 6, pp. 1226–1229, 1995.

[8] R. Alert and X. Trepat, “Physical Models of Collective Cell Migration,” Annu. Rev. Condens. Matter Phys., vol. 11, pp. 77–101, 2020.

[9] T. E. Angelini, E. Hannezo, X. Trepat, M. Marquez, J. J. Fredberg, and D. A. Weitz, “Glasslike dynamics of collective cell migration,” Proc. Natl. Acad. Sci. U. S. A., vol. 108, no. 12, pp. 4714–4719, 2011.

[10] S. Z. Lin, W. Y. Zhang, D. Bi, B. Li, and X. Q. Feng, “Energetics of mesoscale cell turbulence in two-dimensional monolayers,” Commun. Phys., vol. 4, no. 21, pp. 1–9, 2021.

[11] H. H. Wensink et al., “Meso-scale turbulence in living fluids,” Proc. Natl. Acad. Sci. U. S. A., vol. 109, no. 36, pp. 14308–14313, 2012.

[12] A. Doostmohammadi, J. Ignés-Mullol, J. M. Yeomans, and F. Sagués, “Active nematics,” Nat. Commun., vol. 9, no. 1, 2018.

[13] T. B. Saw, W. Xi, B. Ladoux, and C. T. Lim, “Biological Tissues as Active Nematic Liquid Crystals,” Adv. Mater., vol. 30, no. 47, pp. 1–12, 2018.

[14] A. Ardaseva and A. Doostmohammadi, “Topological defects in biological matter,” Nat. Rev. Phys., 2022.

[15] B. Fardin, M.A.; Ladoux, “Living proof of effective defects,” Nat. Phys., vol. 17, no. 2, pp. 172–173, 2021.

[16] T. B. Saw et al., “Topological defects in epithelia govern cell death and extrusion,” Nature, vol. 544, no. 7649, pp. 212–216, 2017.

[17] K. Kawaguchi, R. Kageyama, and M. Sano, “Topological defects control collective dynamics in neural progenitor cell cultures,” Nature, vol. 545, no. 7654, pp. 327–331, 2017.

[18] L. A. Solchaga, K. J. Penick, and J. F. Welter, “Chondrogenic differentiation of bone marrow-derived mesenchymal stem cells: tips and tricks,” Methods Mol. Biol., vol. 698, no. 12. 2011.

[19] G. Duclos, C. Erlenkämper, J. F. Joanny, and P. Silberzan, “Topological defects in confined populations of spindle-shaped cells,” Nat. Phys., vol. 13, no. 1, pp. 58–62, 2017.

[20] V. M. Worlitzer, G. Ariel, A. Be’er, H. Stark, M. Bär, and S. Heidenreich, “Turbulence-induced clustering in compressible active fluids,” Soft Matter, vol. 17, no. 46, pp. 10447–10457, 2021.

[21] Y. Sasano, I. Mizoguchi, M. Kagayama, L. Shum, P. Bringas, and H. C. Slavkin, “Distribution of type I collagen, type II collagen and PNA binding glycoconjugates during chondrogenesis of three distinct embryonic cartilages,” Anat. Embryol. (Berl)., vol. 186, no. 3, pp. 205–213, 1992.

[22] J. H. Hong et al., “TAZ, a transcriptional modulator of mesenchymal stem cell differentiation,” Science, vol. 309, no. 5737, pp. 1074–1078, 2005.

[23] I. Sekiya, B. L. Larson, J. R. Smith, R. Pochampally, J. Cui, and D. J. Prockop, “Expansion of Human Adult Stem Cells from Bone Marrow Stroma: Conditions that Maximize the Yields of Early Progenitors and Evaluate Their Quality,” Stem Cells, vol. 20, no. 6, pp. 530–541, 2002.

[24] A.-A. A. M. Faten and A. A. Zaki, “The impact of confluence on BMMSC proliferation and osteogenic differentiation,” Int. J. Hematol. Stem Cell Res., vol. 11, no. 2, pp. 121–132, 2016.

[25] Z. Zhou et al., “Machine learning forecasting of active nematics,” Soft Matter, vol. 17, no. 3, pp. 738–747, 2021.

[26] T. Turiv et al., “Topology control of human fibroblast cells monolayer by liquid crystal elastomer,” Sci. Adv., vol. 6, no. 20, pp. 1–11, 2020.

[27] B. Yu, X. Zhang, and X. Li, “Exosomes derived from mesenchymal stem cells,” Int. J. Mol. Sci., vol. 15, no. 3, pp. 4142–4157, 2014.

[28] S. Preibisch, S. Saalfeld, and P. Tomancak, “Globally optimal stitching of tiled 3D microscopic image acquisitions,” Bioinformatics, vol. 25, no. 11, pp. 1463–1465, 2009.

[29] K. Zuiderveld, Contrast Limited Adaptive Histogram Equalization. Academic Press, Inc., 1994.

[30] R. Rezakhaniha et al., “Experimental investigation of collagen waviness and orientation in the arterial adventitia using confocal laser scanning microscopy,” Biomech. Model. Mechanobiol., vol. 11, no. 3–4, pp. 461–473, 2012.

[31] D. A. Rennerfeldt, J. S. Raminhos, S. M. Leff, P. Manning, and K. J. Van Vliet, “Emergent heterogeneity in putative mesenchymal stem cell colonies: Single-cell time lapsed analysis,” PLoS ONE, vol. 14, no. 4. 2019.

